# Improving the detection of *A. cantonensis* in brain tissues of mammalian hosts

**DOI:** 10.1101/2022.12.07.518866

**Authors:** Micaela Arango, Sofía Delgado-Serra, Lee Haines, Claudia Paredes-Esquivel

## Abstract

*Angiostrongylus cantonensis* is an invasive nematode parasite that can cause eosinophilic meningitis in many vertebrate hosts, including humans. This parasite is spreading rapidly through the six continents, with Europe being the final frontier. Sentinel surveillance may be a cost-effective surveillance strategy to monitor the arrival of this pathogen to new geographical regions as this can be easily expanded to incorporate symptomatic wildlife. Necropsy and tissue digestion techniques are often used to recover helminth parasites from vertebrate host tissues, however their application to detect brain parasites is poorly utilized. Here we show that employing these techniques in combination can 1) help resolve false positive and negative animals, 2) provide accurate parasitic load values and 3) establish an accurate prevalence of *A. cantonensis*. Our adapted tissue digestion technique can be easily performed, especially in wildlife hospitals where animal infections often precede human cases. Early detection of the parasite increases the efficacy of prevention, treatment, and disease control strategies not only in humans, but also in susceptible animal populations.

**Key Findings:**

- Optimized brain digestion techniques can detect parasitic helminths (*A. cantonensis*) in mammalian hosts.
- Accuracy identifying neurotropic parasitic infections can be increased if a standard digestion protocol is applied.
- The technique presented here can be easily implemented to detect brain nematodes in wildlife hospitals.

## Introduction

Several neurotropic nematodes are now considered globally emerging (Sorvillo *et al*., 2002; Walker and Zunt, 2005). Among these, *Angiostrongylus cantonensis*, the main cause of zoonotic eosinophilic meningitis worldwide, is spreading at an alarming rate (McAuliffe *et al*., 2019). Almost 3000 registered cases caused by this nematode have been reported worldwide, which has led to its designation as an emerging zoonosis (Wang *et al*., 2012). Paralysis, astasia, lateral recumbency, and bicycling movement (Delgado-Serra *et al*., 2022) have been described as the most common clinical manifestations of neuroangiostrongyliasis (NA).

In Europe, the parasite was reported recently in Mallorca, Spain (Paredes-Esquivel *et al*., 2019) and since then, active surveillance has been proposed as a priority in continental Europe (Gonzálvez and Ruiz de Ybáñez, 2022). We recently suggested using hedgehogs for sentinel surveillance as this is a cost-effective strategy for its early detection (Delgado-Serra *et al*., 2022). Symptomatic wildlife species may be used as sentinels, as these show typical clinical manifestations that may alert veterinarians on parasite circulation in other regions. In Australia, where the disease is endemic, dogs, tawny frogmouths and brushtail possums have been proposed as sentinel hosts (Lunn *et al*., 2012; Ma *et al*., 2013).

Necropsies, followed by enzymatic digestion of tissues, are considered easy and cost-effective techniques to detect parasites in host organs as neither protocol requires expensive reagents, specialized machinery or trained staff. Tissue digestion is not a new methodology and has proved valuable for isolating protozoan and helminth parasites from several host tissues (Dubey, 1998b, 1998a). Members of the *Trichinella* genus are routinely isolated from meat samples using magnetic stirrer-assisted digestion (Mayer-Scholl *et al*., 2017). Nevertheless, digestion procedures are less commonly used for neurotropic parasites; it is notoriously difficult to isolate brain-localised parasites as the host’s skull must be opened to assess the tissue. The brain is the primary tissue for neurotropic helminths to develop (several life-stages of the same parasite can be present in different cerebral areas at the same time). Some nematodes also show tissue specificity for a specific brain microenvironment, which can further complicate isolation (Janecek *et al*., 2014; Spratt, 2015).

During our initial surveillance in hedgehogs, we noticed that some animals showed clinical manifestations compatible with NA, yet, surprisingly upon initial investigation, these animals tested negative. This led us to design a digestion technique to definitively confirm infection status. We evaluated the efficacy of using a combination of necropsy followed by enzymatic digestion of brain tissue to detect *A. cantonensis* in sentinel hosts. We expect this modified technique will be useful for other helminths infecting the mammalian brain.

## Materials and methods

### Methodology

We designed a timesaving, tissue-clearing digestion technique by adapting and optimizing several published protocols (Dubey, 1998b; Mayer-Scholl *et al*., 2017; Waindok *et al*., 2019) for other organ tissues. As helminths are sensitive to HCl (Dubey, 1998a), incubation times are key for retrieving non-damaged specimens, and why a shorter and non-destructive protocol was needed so to preserve worms’ integrity for identifying characteristics (Dubey, 1998a, 1998b). The digestion solution was prepared with distilled water instead of saline as some nematode larvae are susceptible to salt concentration (Dubey, 1998a). A 1% pepsin concentration was used in alignment with most other published protocols (Table 1). To mimic host body temperature, digestion solution preparation was performed at 37 °C.

**Table 1.** Summary of enzymatic digestion techniques used for isolating nematodes from brain tissues.

| Nematode | Host | Enzyme concentration | HCl concentration | Solvent | Time | Reference |
| --- | --- | --- | --- | --- | --- | --- |
| *Toxocara canis* and *Toxocara cati* | Mouse | 1% (w/v) pepsin | 300 mM HCl | Distilled water | 2 h | (Janecek *et al.*, 2014) |
| *Haelicephalobus gingivalis* | Horse | 1% (w/v) pepsin | 1mL HCl* | Distilled water | 48 – 72 h | (Pintore *et al.*, 2017) |
| *Aelurostrongylus abstrusus* | Mouse | 0.3 % (w/v) pepsin | HCl*/ ** | Distilled water | 75 min. | (Colella *et al.*, 2019) |
| *Baylisascaris procyonis* | Racoon | 1% (w/v) pepsin | 1% HCl* | 0.85% (w/v) Saline | 2 – 2.5 h | (Kazacos, 2008) |
| *Angiostongylus cantonensis* | Hedgehog | 1% (w/v) pepsin | 300 mM HCl | Distilled water | 15-20 min. | This study |

All digestions were conducted at 37 °C, \* information on the initial concentration of this reagent was not provided by the author. \*\*("100 mL HCl Solution pH: 2.2")

Prior to dissecting suspected parasite-infected tissue, it is important to:

- Record specimen information: identification number, date of necropsy, date of enzymatic digestion processing.
- Defrost head samples overnight (4 °C in the fridge).
- Freshly prepare 100 ml of pepsin solution using 300 mM HCl (LabKem) and 1 % pepsin (DINKO instruments).
- Use a stir bar to gently mix components while heating at 37 °C for 1h.

#### Procedure

1. Using a basic disposable surgical scalpel, make two incisions (~15 mm); the first one from the frontal bone sides (near the eye-section) to the parietal bone (the back of the head, where the vertebral column starts); then make a second transversal incision across the anterior end of frontal bones (Figure 1). Using tweezers, gently separate the braincase from the brain*.
2. After stereomicroscopic examination of the removed braincase, the brain itself and the specimen’s head for parasites, place the brain (average hedgehog brain weight: 1.42 g) into a 100 mL glass beaker.
3. Add 25** ml of the pre-warmed 1% pepsin solution and place on a heated stir plate.
4. Keep stirring the sample + Pepsin solution at 37°C for 15-20 minutes at 400 rpm.
5. Place digested tissue into glass petri dishes and observe with a stereomicroscope (5x). * If possible, pre-digestion, try to separate the brain into different parts (Figure 1). ** For larger brain sizes, the volume of pepsin solution must be increased in a ratio of 1:16 (1 g of tissue/16 mL of solution).

**Figure 1.**
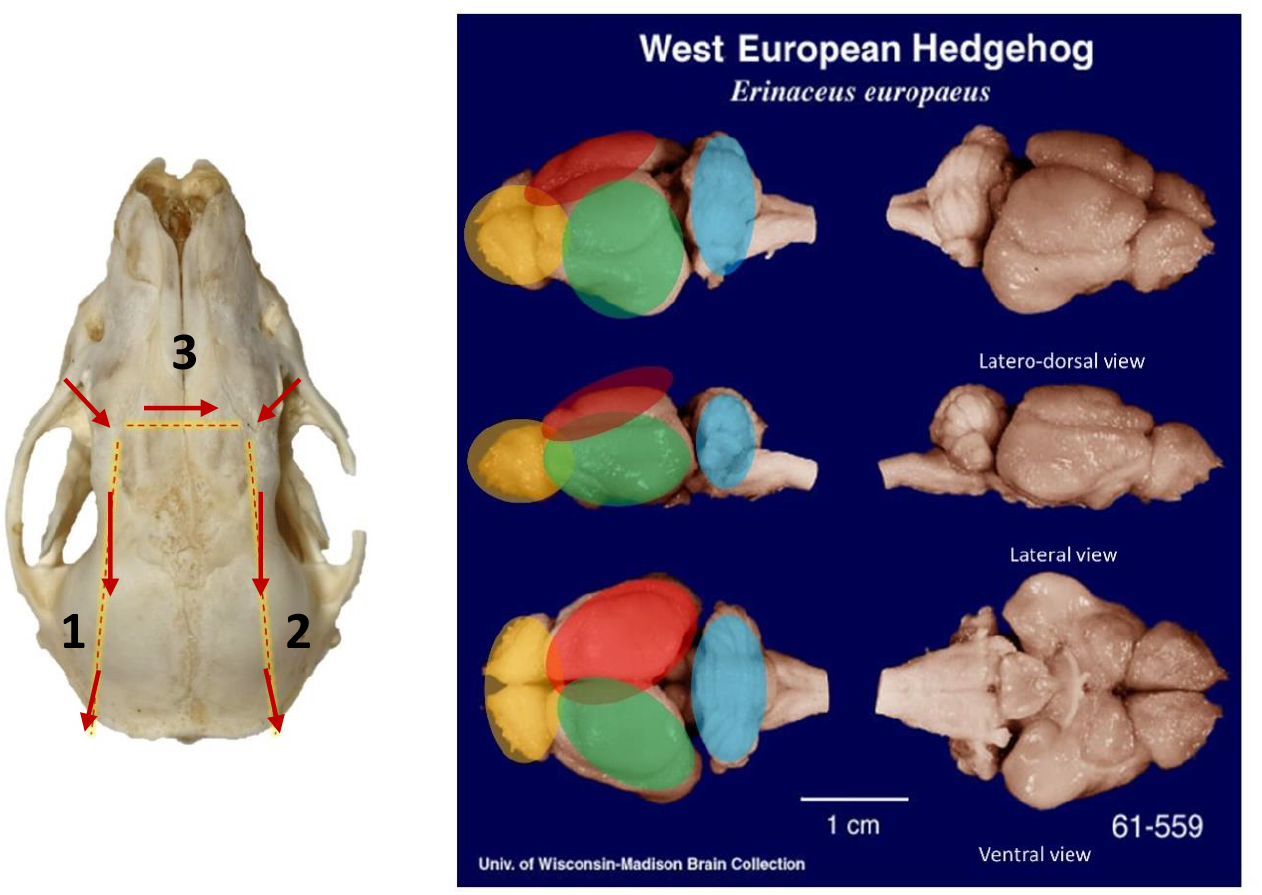
Scheme of the skull and brain of an erinaceid. Left: hedgehog skull showing the incisions and direction made for the brain extraction (modified from http://skullbase.info/). Right: regions of the hedgehog brain: in green, cerebrum left hemisphere; in red, cerebrum right hemisphere; in light blue, cerebellum and in yellow, olfactory bulbs (modified from (Comparative Mammalian Brain Collections, brainmuseum.org).

## Results

The adapted protocol for digesting brain tissue is an effective method to recover obscured helminths (Figure 2). In most cases (7 out of 9), we were able to recover parasites hidden deep within the brain parenchyma. These parasites were impossible to detect during ocular inspection at necropsy and thus the animal had been incorrectly classified as NA negative. A total of 92 nematodes were recovered directly by visual inspection after necropsy, either in tissues from the subarachnoid space or in the skull, while a further 58% (127 nematodes) were liberated from the parenchyma only after tissue digestion. The average parasite burden per hedgehog using this technique increased from 10.2 (pre-digestion) to 14.1 post-digestion. By digesting brain tissue, we can confidently report eight of the nine hedgehogs were positive for *A. cantonensis*. One hedgehog (AaAL1) tentatively diagnosed negative to NA was reclassified as positive as no parasites were observed during necropsy but were evidenced after digestion. Of all the nematodes recovered after digestion, we calculated 57.9 % of *A. cantonensis* specimens remain undetected if diagnosis relies solely on necropsy. Interestingly, in the case of hedgehog AaPA2 post-digestion (Figure 3), 48 additional parasites were recovered; 19.2 % of the specimens remained unidentified in terms of sex/life cycle stage as they were damaged during nematode collection. Five of the nine hedgehogs positive for *A. cantonensis* were at their juvenile state, whereas the rest were classified as adults.

**Figure 2.**
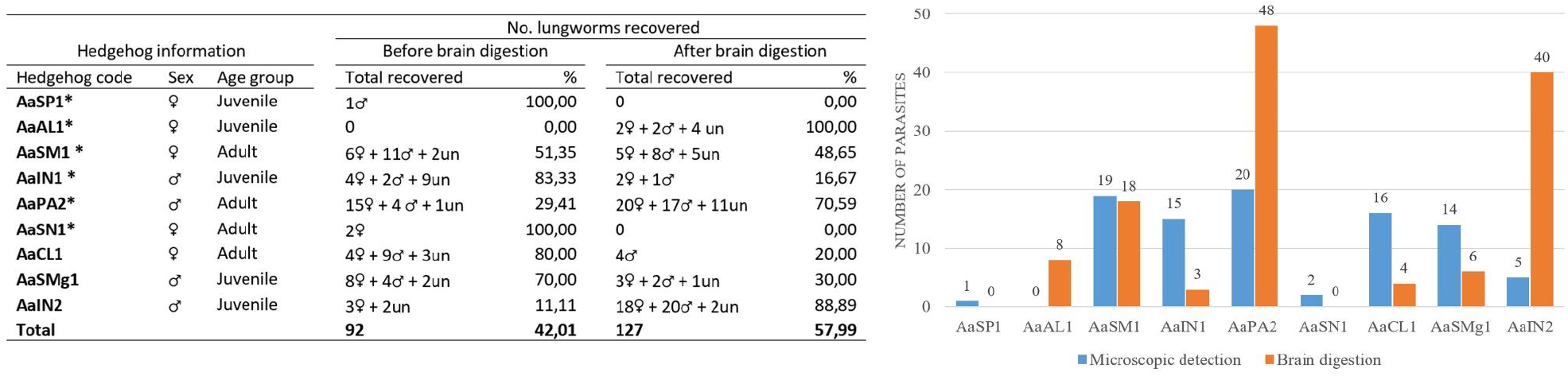
Summary of *A. cantonensis* nematodes recovered before and after tissue digestion. On the left, a table showing the recovery of *A. cantonensis* for each hedgehog, showing nematode sex and the total worm burden per hedgehog. On the right, a bar chart highlights the total nematode load before (blue) and after (orange) brain digestion by plotting the number of isolated parasites (y-axis) per hedgehog (x-axis). un: sex unidentified /broken specimen.

**Figure 3.**
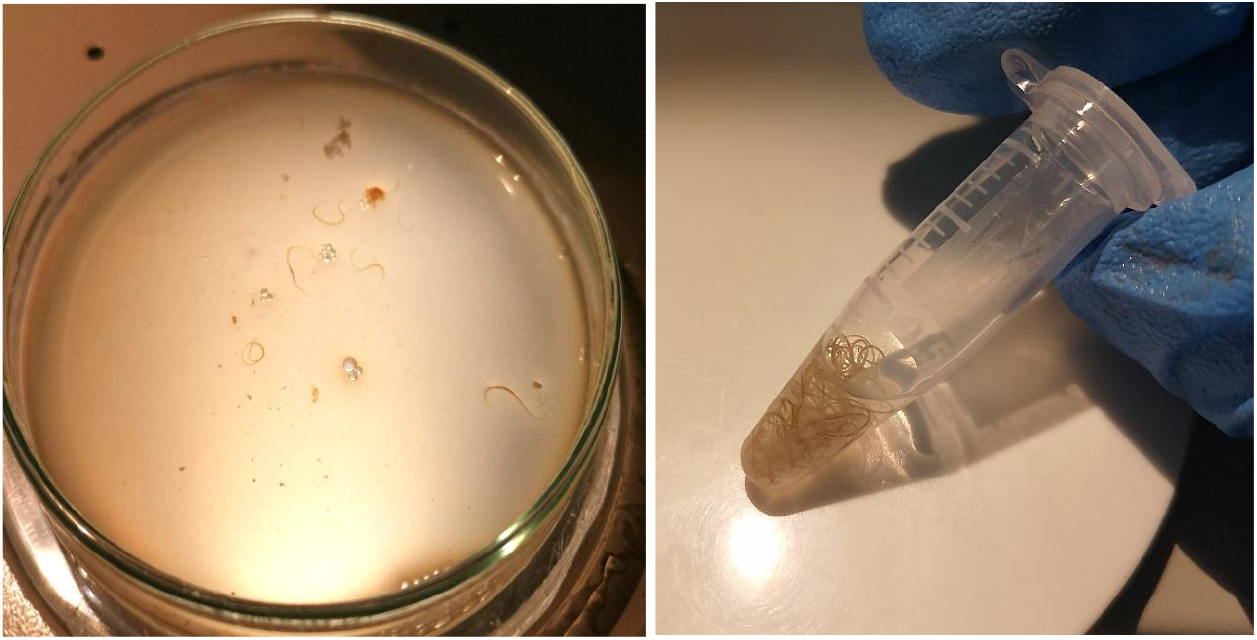
Isolated *A. cantonensis* post pepsin digestion. Left panel: a petri dish containing *A. cantonensis* worms after digesting the brain of a single hedgehog. Right panel: collection of isolated worms that showcase the signature barber pole morphology of *A. cantonensis*.

## Discussion

Enzyme-based digestion techniques have previously been used to isolate *A. cantonensis* from intermediate (Jarvi *et al*., 2012, 2015) and paratenic host tissues (Graeff-Teixeira and Morera, 1995). Although digestion-based methods are also expected to work on definitive and accidental host tissues, further experiments confirming this remain outstanding. To the best of our knowledge, digestion techniques with brain tissues have only been used for the following neurotropic nematodes*: Baylisascaris procyonis* (Kazacos, 2008), *Toxocara* spp. (Janecek *et al*., 2014), *Haelicephalobus gingivalis* (Pintore *et al*., 2017), *Aelurostrongylus abstrusus* (Colella *et al*., 2019) (Table 1). The latter protocol was originally designed for snail muscles and not optimized for the softer brain tissues (Colella *et al*., 2016), which could have resulted in parasite degradation. With the exception the study by Janecek *et al* (2014), the remaining studies did not provide enough information on reagents concentration, particularly the original concentration of HCl, impairing reproducibility. Parasite integrity, enzyme concentration and efficacy are highly sensitive to salt, pH and length of incubation at temperature (Dubey, 1998b, 1998a). The optimization of an affordable, reliable, reproducible and rapid digestion protocol as we present here is a desirable tool for intact neurotropic-helminth recovery. Although we report some of the isolated worms were damaged or broken, this is likely the result of freezing and/or dissection artefacts rather than a result of enzymatic digestion as previously reported (Sánchez-Alonso *et al*., 2021). Also, in accidental hosts, some worms are expected to degenerate as most will not be able to reach the lungs (Prociv and Turner, 2018). Working with frozen carcasses is not optimal; it is more of a practical step in wildlife hospitals where animals can be safely stored until staff have enough time to perform the labor-intensive necropsies. Despite these limitations, most nematodes we isolated were undamaged. Intact parasite isolation is the key for many biomedical and epidemiological studies (Barratt *et al*., 2016). For *A. cantonensis*, the infection routes have recently been under scrutiny as this information is essential for successful human therapeutic decisions; better prognosis are seen when treatment is applied at the beginning of the sub-arachnoid phase (Prociv and Turner, 2018). This stage characterized by the migration of the worms into the subarachnoid space has not been characterized in humans and animal hosts are often used as models.

All necropsied hedgehogs were infected with adult worms (table 2) which may indicate that mortality occurs at least 11-13 days after infection, if the life cycle in rats is used as reference (Mackerras and Sandars, 1954). The detection of adult worms within the parenchyma may also indicate that subarachnoid phase is likely to be followed by the migration of the nematodes to the deeper tissues of the hedgehog brain. However, this may be just, an artifact of postmortem migration since nematodes are known to invade the surrounding tissues of their hosts after death (Prociv and Turner, 2018). More studies are needed to clarify the trajectory of the rat lungworm in hedgehogs as their role as definitive hosts have been previously discussed (Paredes-Esquivel *et al*. 2019).

Although tissue biopsies are routinely taken to diagnose cases of human NA (Morton *et al*. 2013), tissue digestion techniques are not used on human brain tissue. Furthermore, for human samples, there is little information on the efficacy of tissue digestion protocols for detecting nematode infections, despite several reports on the use of these techniques for the identification of viruses (Beck *et al*. 2004) and bacteria from gastric, esophageal, and colorectal biopsies (Zhang *et al*. 2019). Consequently, this methodology remains open for investigations looking for nematodes embedded in human tissue.

Previous studies on the patterns of parasite dispersal using mathematical modeling show that Europe and other currently non-endemic regions will increase in suitability for the establishment *A. cantonensis* as oscillations in humidity, precipitation and temperature levels could directly affect the life cycle of the parasite and therefore its transmission (York, Butler and Lord, 2014). Creating robust and simple diagnostic tools will help detect these emerging parasites before numbers of human cases increase or susceptible fauna become at higher risk of infection.

By detecting emerging parasites early, disease control interventions could be pre-emptively deployed to prevent further spreading opportunities, a more accurate idea of epidemiological status can be drawn, and the overall risk for humans, their companion animals, and livestock can be accurately assessed (Morgan *et al*., 2012). Here we present a method to digest post-mortem brain tissue using a modified digestion protocol that takes only 20 minutes. This protocol would enhance current animal examinations and allow for earlier detection of *A. cantonensis* in veterinary hospitals where symptomatic animals with NA are concentrated. Our adapted protocol is a faster and more efficient way to accurately determine disease prevalence and parasite load in post-mortem exams and could likely be useful for detecting other tissue burrowing parasitic nematodes in different mammalian species.

## Acknowledgements (optional)

We thank the staff at Consorci per a la Recuperació de la Fauna de les Illes Balears for providing frozen hedgehogs diagnosed with NA. We would also like to thank the program SOIB Recerca e Innovació, which aims to integrate young scientists in established research groups from the Universitat de les Illes Balears.

## Author Contributions (mandatory)

CPE and SD conceived and designed the study. MA and SD conducted data gathering. SD, MA, LH and CPE analyzed the data. All authors wrote and edited the article.

## Financial Support (mandatory)

This work has been sponsored by the *Comunitat Autònoma de les Illes Balears* through the *Direcció General de Política Universitària i Recerca* with funds from the Tourist Stay Tax Law (PDR2020/61 – ITS2017-006). Funding for publication in open access was obtained from the *Pla Anual d’Impuls del Turisme Sostenible* de les Illes Balears and the FEDER funds. Michaela Arango’s contract was funded by SOIB Recerca e Innovació.

## Ethical Standards

Not applicable here

